# Towards reliable reconstruction of the mouse brain thalamocortical connectivity using diffusion MRI

**DOI:** 10.1101/2022.12.28.522151

**Authors:** Tanzil Mahmud Arefin, Choong Heon Lee, Zifei Liang, Harikrishna Rallapalli, Youssef Z. Wadghiri, Daniel H. Turnbull, Jiangyang Zhang

## Abstract

Diffusion magnetic resonance imaging (dMRI) tractography has yielded intriguing insights into brain circuits and their relationship to behavior in response to gene mutations or neurological diseases across a number of species. Still, existing tractography approaches suffer from limited sensitivity and specificity, leading to uncertain interpretation of the reconstructed connections. Hence, in this study, we aimed to optimize the imaging and computational pipeline for reliable reconstruction of the mouse brain thalamocortical network. We developed a dMRI-based atlas of the mouse forebrain with structural labels imported from the Allen Mouse Brain Atlas (AMBA). Using the atlas and tracer data from the Allen Mouse Brain Connectivity Atlas (AMBCA) as ground truth, we investigated the accuracy of reconstructed node-to-node thalamocortical structural connectivity and effects of imaging and tractography parameters. Our results suggest that these parameters significantly affect tractography outcomes and our atlas can be used to investigate macroscopic structural connectivity in the mouse brain. Furthermore, tractography in mouse brain gray matter still face challenges and need improved imaging and tractography methods.

## 1. Introduction

Understanding the structural and functional networks in the brain is one of the major areas of neuroscience research. Over the past decade, tremendous efforts have been made in mapping neural architecture at various scales, from tract tracing at the mesoscopic scale (Hunnicutt et al., 2014; Oh et al., 2014; Zingg et al., 2014) to the identification of cellular-level connections and the synaptic molecular properties (Siegle et al., 2021; Winnubst et al., 2019). Mapping of neural networks has also been reported in multiple species, ranging from C. elegans to human (Cook et al., 2019; Majka et al., 2020; Oh et al., 2014; Scheffer et al., 2020; Szczupak et al., 2021; Xu et al., 2021).

Structural connectivity in the mouse brain has been a main focus due to the large number of mouse models of neurological diseases together with accessibility of mouse genomic sequence information and a large number of publicly available resources on the mouse brain (Sandberg et al., 2000; Siddiqui et al., 2005; Zapala et al., 2005). A leading example is the Allen Mouse Brain Connectivity Atlas (AMBCA) (http://connectivity.brain-map.org), a systematic analysis of long-distance axonal connectivity using recombinant adeno-associated virus (AAV) expressing enhanced green fluorescent protein (EGFP) as the tracer. AMBCA presents a mesoscale connectome of the mouse brain (Kuan et al., 2015; Oh et al., 2014), which is also integrated with genome-scale collection of cellular resolution gene expression profiles (Goldowitz, 2010; Lein et al., 2007; Ng et al., 2009). This and similar resources from other groups (Hunnicutt et al., 2014; Jeong et al., 2016; Zingg et al., 2014) have led to greater understanding of the mouse brain mesoscopic connectivity and its organization principles (Coletta et al., 2020; Rubinov et al., 2015).

Although the uses of chemical and viral tracers have allowed direct visualization of neural connections, one limitation of the technique is the inability to examine a large number of circuitries within a single brain. In contrast, diffusion magnetic resonance imaging (dMRI) tractography (Jbabdi et al., 2015; Mori and van Zijl, 2002) permits non-invasive mapping of multiple pathways in the entire brain. Combining advanced dMRI tractography and computational tools further allows non-invasive exploration of structural connectivity in rodent brains and alterations due to genetic modifications or pathological conditions (Arefin et al., 2021; Arefin et al., 2017; Calabrese et al., 2015; Degiorgis et al., 2022; Mechling et al., 2016; Moldrich et al., 2010; Ren et al., 2007; Wang et al., 2020a; Wang et al., 2006; White et al., 2020; Wu et al., 2022; Yee et al., 2018). However, more wide-spread applications of dMRI tractography in connectome analysis has been hindered by its relatively low sensitivity and specificity, especially in gray matter (GM) structures, (Aydogan et al., 2018; Calamante, 2019; Schilling et al., 2019; Thomas et al., 2014) due to the lack of direct links between dMRI signals and the underlying cellular structures (e.g. axons) and the low spatial resolution limitation of MRI. A recent study by Trinkle *et al*. (Trinkle et al., 2021) compared connectomes based on AMBCA and dMRI data and revealed considerable differences between them, in particular, the lack of connections in the thalamocortical network. However, in depth quantification of the thalamocortical circuitry that can be reconstructed using dMRI tractography as compared to the corresponding AMBCA is lacking. For the general use of dMRI to examine connectome in normal or diseased mouse brains, it is important to obtain detailed knowledge on its sensitivity and specificity and locate optimal parameters for dMRI tractography.

For systematic analysis of tractography results together with resources such as AMBCA, an MRI-based atlas with structural labels that are compatible with the Allen Mouse Brain Atlas (AMBA) is necessary. To date, a number of MRI-based mouse brain atlases and datasets have been reported, covering both adult and developing mouse brains (Ali et al., 2005; Badea et al., 2007a; Badea et al., 2007b; Bock et al., 2006; Chen et al., 2006; Chuang et al., 2011; Dorr et al., 2007; Dorr et al., 2008; Johnson et al., 2007; Kovacevic et al., 2005; Lee et al., 2005; Ma et al., 2005; MacKenzie-Graham et al., 2004; Richards et al., 2011; Sharief et al., 2008; Steadman et al., 2014; Szulc et al., 2015; Ullmann et al., 2013). A summary of these atlases and datasets is provided in supplementary material table S1. Many mouse brain atlases were created based on *ex vivo* T_1_/ T_2_ MRI with high spatial resolution (~ 30 um) and strong tissue contrasts in the adult mouse brain, mainly reflecting tissue water and myelin contents. Atlases based on dMRI and other MRI contrasts have emerged over the past decade (Aggarwal et al., 2009; Calabrese et al., 2015; Chuang et al., 2011; Jiang and Johnson, 2011; Szulc et al., 2015). Compared to T_1_/T_2_ MRI, dMRI provides additional contrasts reflecting tissue microstructural organization (Mori and Zhang, 2006) and can be used to map macroscopic connectivity in the mouse brain (Aydogan et al., 2018; Calabrese et al., 2015; Keifer et al., 2015). The existing atlases, however, often do not have detailed cortical and subcortical structural labels, mainly because MRI data lack the resolution and tissue contrasts comparable to histology for delineation of subregions in GM such as deep thalamic nuclei or cortical layers.

To meet this demand, we have constructed a dMRI based mouse brain atlas by importing the AMBA structural labels in our previously published group average diffusion tensor images of the adult mouse brain (Chuang et al., 2011) and constructed a mapping between the DTI atlas and AMBA. The mapping of AMBA data to our atlas provided structural labels with high granularity and allowed us to examine mouse brain tractography results with viral tracer results in AMBCA in the same space consistently. Hence, we were able to use the node specific tracer maps as the gold standard to quantify the accuracy of tractography-based reconstruction of the thalamocortical connections in ex vivo dMRI data acquired from a separate group of mouse brains.

## 2. Materials and Methods

All experimental procedures were approved by the Animal Use and Care Committee at the New York University Grossman School of Medicine.

### 2.1 Animals

Adult C57BL/6J mice (males, P60-P90, n = 16) were used in this study. Six of them were used for in vivo manganese-enhanced MRI (section 2.3). The remaining ten mice were perfused transcardially with 4% paraformaldehyde in 0.1M phosphate buffer saline (PBS) and stored in 4% paraformaldehyde solution for 24 hours before transferred into PBS. Mouse brains kept in the skull were placed in 10 mL syringes filled with Fomblin (Perfluoropolyether, Solvay Solexis Inc.,Thorofare, N), a non-hydrogen containing liquid for susceptibility matching and to prevent dehydration. Detailed procedures on how to prepare the *ex vivo* brain sample can be found in (Arefin et al., 2021).

### 2.2 Diffusion-weighted MRI (DW-MRI) data acquisition

#### Cohort 1

data in this cohort were from a previous study (Chuang et al., 2011). *Ex vivo* imaging was performed on an 11.7 Tesla (T) NMR spectrometer (Bruker Biospin, Billerica, MA, USA) as described earlier (Chuang et al., 2011). Briefly, dMRI data was acquired using 3D diffusion-weighted multiple spin echo sequence at an isotropic spatial resolution of 0.125 mm^3^ with further isotropic interpolation to 0.0625 mm by zero-padding in the k-space.

#### Cohort 2

High resolution *ex vivo* dMRI datasets were acquired on a horizontal 7T MR scanner (Bruker Biospin, Billerica, MA, USA). We used a 72-mm conventional circularly polarized birdcage radiofrequency resonator (Bruker Biospin, Ettlingen, Germany) for homogeneous transmission in conjunction with a four-channel receive-only phased array CryoProbe (CRP) anatomically shaped to tightly cover the mouse head (Bruker Biospin, Ettlingen, Germany) for high sensitivity and a modified 3D diffusion-weighted gradient-and-spin-echo (DW-GRASE) sequence (Aggarwal et al., 2010). Three sets of data were obtained with different imaging parameters (Table 1):

**Table 1:** The imaging parameters of the dMRI datasets used in this study. TE and TR are abbreviations of echo time and repetition time, respectively.

| Parameters | Dataset 1 | Dataset 2 | Dataset 3 |
| --- | --- | --- | --- |
| No. of subjects | 10 | 10 | 10 |
| b-value (s/mm<sup>2</sup>) | 2000 | 5000 | 5000 |
| Number of diffusion encoding directions | 30 | 30 | 60 |
| TE/ TR (ms) | 35/400 | 35/400 | 35/400 |
| Isotropic resolution (mm) | 0.1 | 0.1 | 0.1 |
| Acquisition time (h) | ~ 5.3 | ~ 5.3 | ~ 10.6 |

### 2.3 Manganese-enhanced MRI (MEMRI) data acquisition

Solution of manganese chloride (MnCl_2_) tetrahydrate (Sigma-Aldrich-221279) in isotonic saline (30 mM) was injected intraperitoneally to 6 animals 24 hours before MRI at a dose per weight of 0.5 mmol/kg (62.5 mg MnCl_2_ per kg body weight) as described in (Rallapalli et al., 2020). Animals were anesthetized using isofluorane (3% isofluorane for induction and 1–2% isofluorane during MRI mixed with compressed air at a flow rate of 1.5 L/min through a nose cone) and imaged using the horizontal 7T MRI scanner with the same 4-element phased array CRP for high sensitivity. T_1_-weighted images were acquired using a 3D spoiled gradient echo sequence (TE/TR = 3/30 ms, flip angle = 30°; field of view (FOV) = 1.8 × 0.9 × 1.6 cm; matrix size = 180 × 90 × 160 voxels, 0.1 mm isotropic resolution) with a total imaging time of 30 minutes. Individual subject MEMRI data were first rigidly aligned to the dMRI atlas (Chuang et al., 2011), and group average MEMRI data were computed using an iterative procedure as described in (Chuang et al., 2011). The group average MEMRI data were then aligned to the dMRI atlas using affine transformation.

A summary of the experiments carried out in this study has been provided in the supplementary table S2.

### 2.4 dMRI data processing

Signals from non-brain tissue in the dMRI datasets were manually removed using AMIRA (ThermoFisher Scientific, https://www.thermofisher.com) and corrected for ghosting artifacts likely caused by frequency drifts using rigid alignment implemented in DTIStudio (Jiang et al., 2006). For both cohorts of mice, the six elements of diffusion tensor were determined by log-linear fitting via DTIStudio. The tensor was diagonalized to obtain three eigenvalues (λ_1_, λ_2_, λ_3_) and corresponding eigenvectors (v_1_, v_2_, v_3_). Fractional anisotropy (FA) was then calculated from the eigenvalues (Basser and Pierpaoli, 1996; Mori and Zhang, 2006). Finally, average diffusion-weighted images (aDWI) and FA images were normalized and averaged across all subjects of cohort 1 to create a group averaged mouse brain template (aDWI and FA, dimension: 200 × 280 × 128, resolution: 0.0625 mm) for the dMRI-based adult mouse brain atlas (Chuang et al., 2011).

### 2.5 Image registration pipeline for atlas construction

Fig. 1 demonstrates the image registration pipeline used for mapping the ABMA structural labels (common coordinate framework version 3) into the dMRI template space to construct the dMRI based mouse brain atlas. First, group averaged mouse brain template (DWI and FA images – Fig. 1A) and AMBA (Fig. 1B) were co-registered using landmark-based rigid transformation followed by intensity-based affine and large deformation diffeomorphic metric mapping (LDDMM) transformation implemented in DiffeoMap (www.mristudio.org) as shown in Fig. 1C-D. Second, segmentation of major brain structures (e.g., cortex, hippocampus, and cerebellum) were imported from AMBA and manually adjusted to fit corresponding structures in the dMRI data. We assigned unique intensity values to regions occupied by these major brain structures in the AMBA space and dMRI template space to compute a fine-tuned mapping to carry AMBA to the dMRI template space based on these major brain structures using LDDMM. Third, detailed AMBA structural labels were transferred to the group averaged template image (Fig. 1D) and further manually refined slice by slice along the axial orientation forfeiting attention to the other two orientations as well as to the slices preceding and following throughout the segmentation process Fig. 1E. The level of registration accuracy was measured using landmarks manually placed throughout the brain in the AMBA and dMR images resulting in an average error of 0.15±0.08 mm (Fig. 1F).

**Fig. 1:**
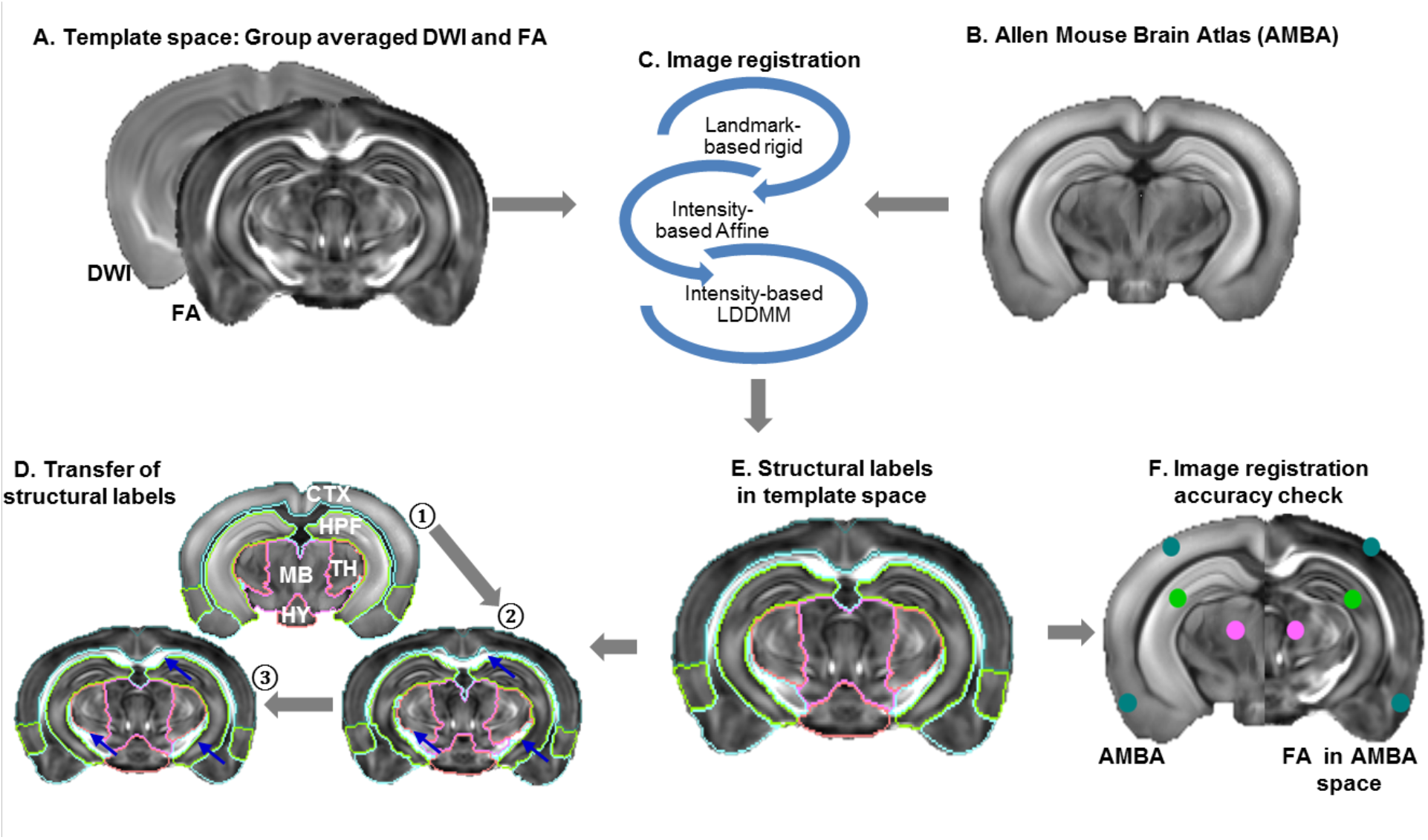
Image registration pipeline for atlas construction: A) Template space: averaged DWI and FA images of the adult mouse brain. B) A matching brain slice from the Allen Mouse Brain atlas (AMBA). C) Coregistration of MR and AMBA images using a series of image registration steps: 1. landmark based rigid transformation, 2. intensity based affine transformation, and 3. intensity based large deformation diffeomorphic metric mapping (LDDMM). D) After co-registration, structural labels of major brain compartments were transferred from AMBA (step ①) to the template space, manually edited (step ②), and a fine-tuned mapping between the template and AMBA was generated based on the labels of major brain compartments (step ③). E) The AMBA structural labels were mapped to the dMRI template using the fine-tuned mapping. F) Assessment of registration accuracy by placing landmarks manually in the template and AMBA images.

### 2.6 Non-constrained and anatomically constrained fiber tractography (NCT and ACT)

dMRI datasets from cohort 2 were analyzed using MRtrix (https://www.mrtrix.org/) (Tournier et al., 2012). *Non-constrained tractography* (*NCT*) was performed using the pipeline described in (Arefin et al., 2021). In brief, using the pre-processed data, fiber orientation distribution (FOD) map was estimated for each subject (Tournier et al., 2007; Tournier et al., 2004), and a whole-brain tractogram was then generated containing a total of 5 million streamlines with minimum fiber length of 3 mm.

For *anatomically-constrained tractography* (*ACT*), in addition to the criteria used for NCT, an image containing five labels of tissue segmentation (5tt) was used to guide the termination and acceptance/rejection criteria during fiber tracking (Smith et al., 2012). The so-called 5tt image file contained cortical GM, sub-cortical GM, white matter (WM), cerebrospinal fluid (CSF), and a null channel ((Supplementary Fig. S1 A). In ACT, any streamlines that entered the cortical GM or sub-cortical region from the WM were counted as valid, and fibers attempting to leave the pre-defined mask were terminated and accepted as valid. Any streamlines attempting to enter CSF or terminating in the WM were truncated and rerouted to find a valid termination point, as known as back-tracking (Fig. 2B-C). A whole-brain tractogram with 5 million streamline was generated using ACT for each subject with minimum fiber length of 3 mm.

Datasets acquired using three different imaging parameters (table 1) and tractography algorithms (NCT or ACT) yielded six conditions (table 2) that were used in the subsequent analysis.

**Table 2:** The six conditions that were compared in this study.

| Conditions | b-values<br>(s/mm<sup>2</sup>) | Number of<br>diffusion directions | Tractography<br>method |
| --- | --- | --- | --- |
| 1<br>(baseline) | 2000 | 30 | NCT |
| 2 | 5000 | 30 | NCT |
| 3 | 5000 | 60 | NCT |
| 4 | 2000 | 30 | ACT |
| 5 | 5000 | 30 | ACT |
| 6 | 5000 | 60 | ACT |

### 2.7 Reconstruction of the thalamocortical connectome and generation of tract density images (TDI)

For each subject, we constructed the thalamocortical connectome using two approaches as following:

#### Open-end tractography

Using 14 cortical regions (supplementary materials table S2) in AMBA, we selected individual fiber projections from the NCT and ACT whole-brain tractograms of each mouse brain.

#### Node-to-node tractography

We used the previous 14 cortical regions and 12 thalamic regions (supplementary materials table S3) in AMBA and selected streamlines from the whole-brain tractogram. Each cortical node served as the seed and a thalamic region as the target region. This approach allowed us to dissect entire circuitry into specific tracts and assess the patterns of fiber projections without superfluous connectivity to other brain regions.

Finally, we generated high resolution tract density images (TDI) for each tractogram as described in (Calamante et al., 2010).

### 2.8 Validation of mouse brain structural connectivity using the tracer maps

From AMBCA, we selected 14 tracer data with injection sites in cortex, identical to the cortical ROIs used for fiber tractography. Detailed information about the injection sites is provided in supplementary table S4. To eliminate the spurious axonal projections, voxels with projection density >0.001 were counted as structurally connected with the injection site (seed region).

As the tractography was performed in subject’s native dMRI space, we transferred each tracer map into each subject’s native dMRI space individually using the image registration steps described earlier and an example is shown in Fig.2. The level of spatial overlaps between the tracer and tractography results was evaluated using the DICE score (Dice, 1945).

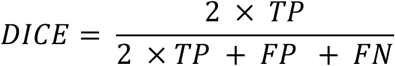

**Fig. 2:**
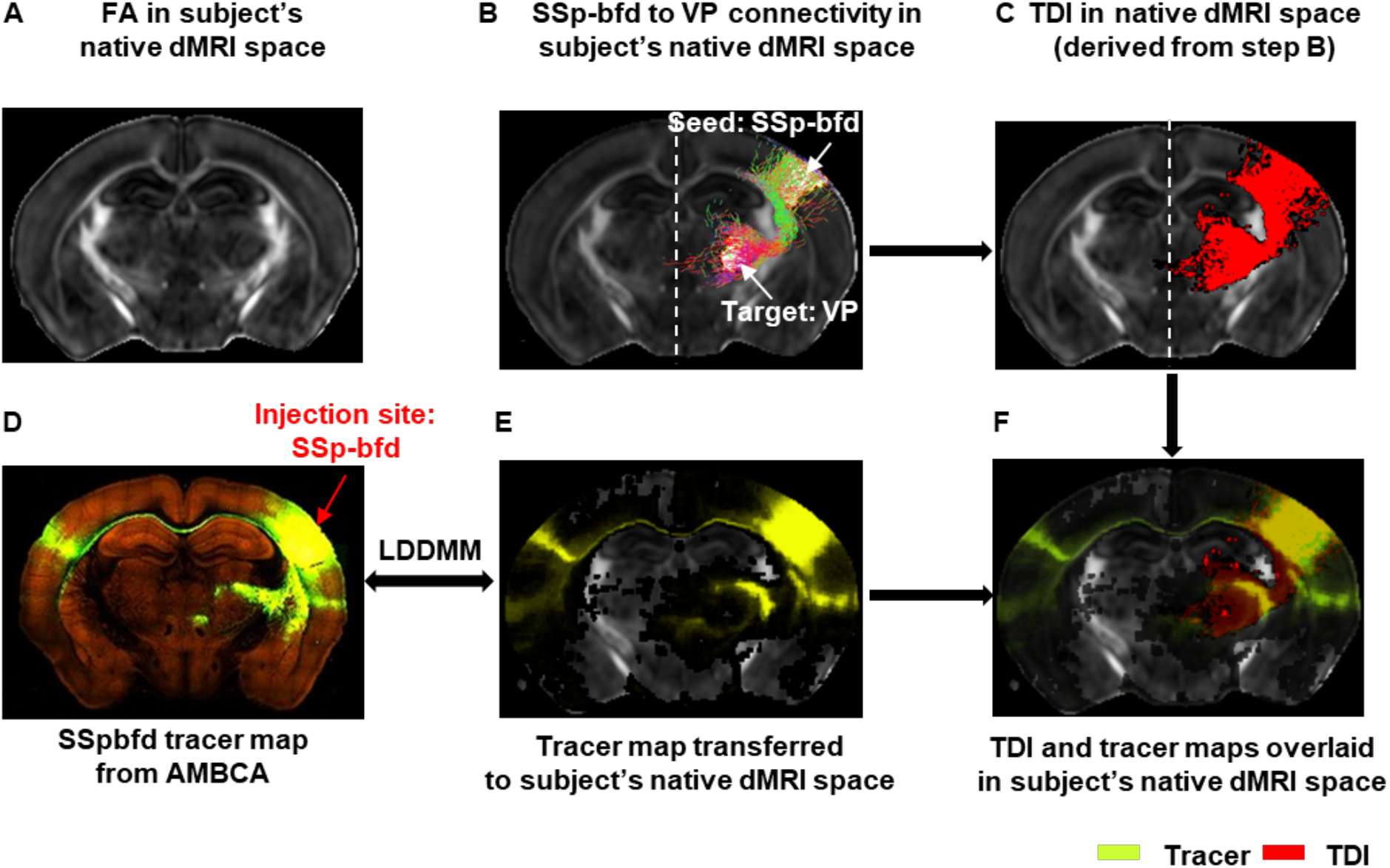
Transferring tracer map to each subject’s native dMRI space for tractography: A) Representative FA image in the native dMRI space. B) Ipsi-lateral tractogram from primary somatosensory barrel field area (SSp-bfd) to ventral posterior complex of the thalamus (VP) extracted from whole brain tractogram shown in the native dMRI space. C) Track density image (TDI) derived from the SSp-bfd to VP fiber tractogram shown in B. D) Tract tracing image from AMBCA – selected injection site: SSp-bfd (ID:112951804). E) Tracer map transferred to the subject’s native dMRI space. F) TDI and tracer maps overlaid on subject’s native dMRI space.

Where TP, FP, FN stand for true positive, false positive, and false negative, respectively. DICE score of 1 represents 100% overlaps/similarities between the estimated thalamocortical connection and the corresponding AMBCA results, while 0 indicates no overlap at all. We defined the level of overlaps in 3 categories: good (DICE > 0.8), moderate (0.6 < DICE < 0.8), and poor (DICE < 0.6).

For all six conditions in table 2, we quantified the DICE scores comparing the TDI and corresponding tracer maps in 2 steps: *1*) *open-end tractography:* whole brain axonal projection maps for 14 cortical regions as described in previous section; *2*) *node-to-thalamus tractography:* extracting tractography streamlines frin each cortical node to the entire thalamus (for example, SSp-bfd to TH, AUDd to TH etc.) from the open-end tractography results. We examined the gross effects of imaging and tractography parameters by computing the DICE scores for each open-end tractograms generated in step 1. Then we computed the DICE for termination points in the thalamus for each cortical node-to-thalamus result (computed in step 2) and categorized the level of similarities between the tractography and tracer maps as good, moderate, or poor based on their spatial overlap in the thalamus using the DICE score. In this way, we were able to identify the conditions that significantly improve the fiber termination in valid target regions and achieve the best match with the corresponding tracer maps for the sub-networks as well as for the specific tracts within the entire thalamocortical network.

### 2.8 Statistical analysis

We used Matlab (www.mathworks.com) and GraphPad Prism (Version 8.4.3 for Windows, GraphPad Software, La Jolla California USA) (www.graphpad.com) for statistical analysis. We performed Two-way ANOVA with Tukey’s multiple comparisons test at α = 0.05 to quantify the level of significance for each of the six conditions.

## 3. Results

### 3.1 dMRI-based adult mouse forebrain atlas

Mapping structural labels from AMBA to the dMRI-based atlas space (Fig. 3) yielded an atlas with a comprehensive set of neuroanatomical regions covering the entire mouse forebrain as mentioned in table 3 and Fig. 4. Detailed description of the structures can be found in the supplementary material.

**Fig. 3:**
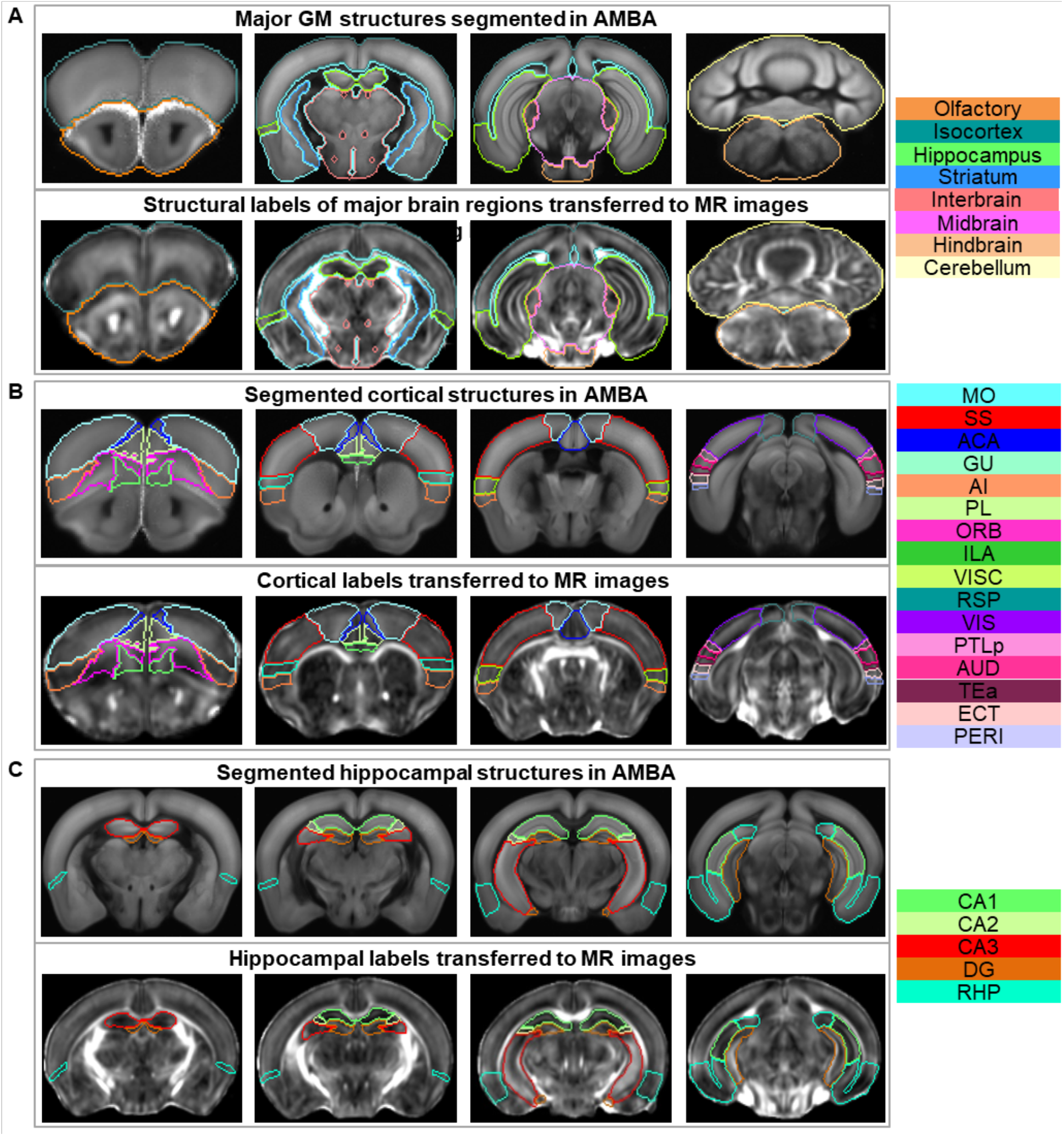
AMBA structural labels transferred to the dMRI space: A) Major GM structures. B) Cortical structures. C) Hippocampal structures. Upper and lower panels of A, B, and C show the structural labels in AMBA space and in dMRI space after coregistration respectively. The rightmost column lists the color-coded regions included in the corresponding atlas. Abbreviations of the anatomical structures are same as defined in the AMBA (https://mouse.brain-map.org/static/atlas).

**Table 3:** Names and number of structures of the dMRI-based atlas.

| Parent structures | Name of the atlases | No. of structures |  |
| --- | --- | --- | --- |
|  |  | Left | Right |
| (a) Cerebrum | Whole brain (WB) | 11 | 11 |
| (b) Cortical plate | Olfactory area (OLF) | 10 | 10 |
|  | Isocortex (CTX) – only major regions | 17 | 17 |
|  | Isocortex (CTX) – with subregions | 40 | 40 |
|  | Hippocampus (HPF) – only major regions | 5 | 5 |
|  | Hippocampus (HPF) – with subregions | 9 | 9 |
| (c) Cortical subplate | Cortical subplate (CTXsp) | 6 | 6 |
| (d) Cerebral nuclei | Striatum (STR) | 4 | 4 |
|  | Pallidum (PAL) | 4 | 4 |
| (e) Brain stem | Interbrain – Thalamus (TH) | 16 | 16 |
|  | Interbrain – Hypothalamus (HY) | 4 | 4 |
|  | Midbrain (MB) | 3 | 3 |
|  | Hindbrain (HB) | 6 | 6 |

**Fig. 4:**
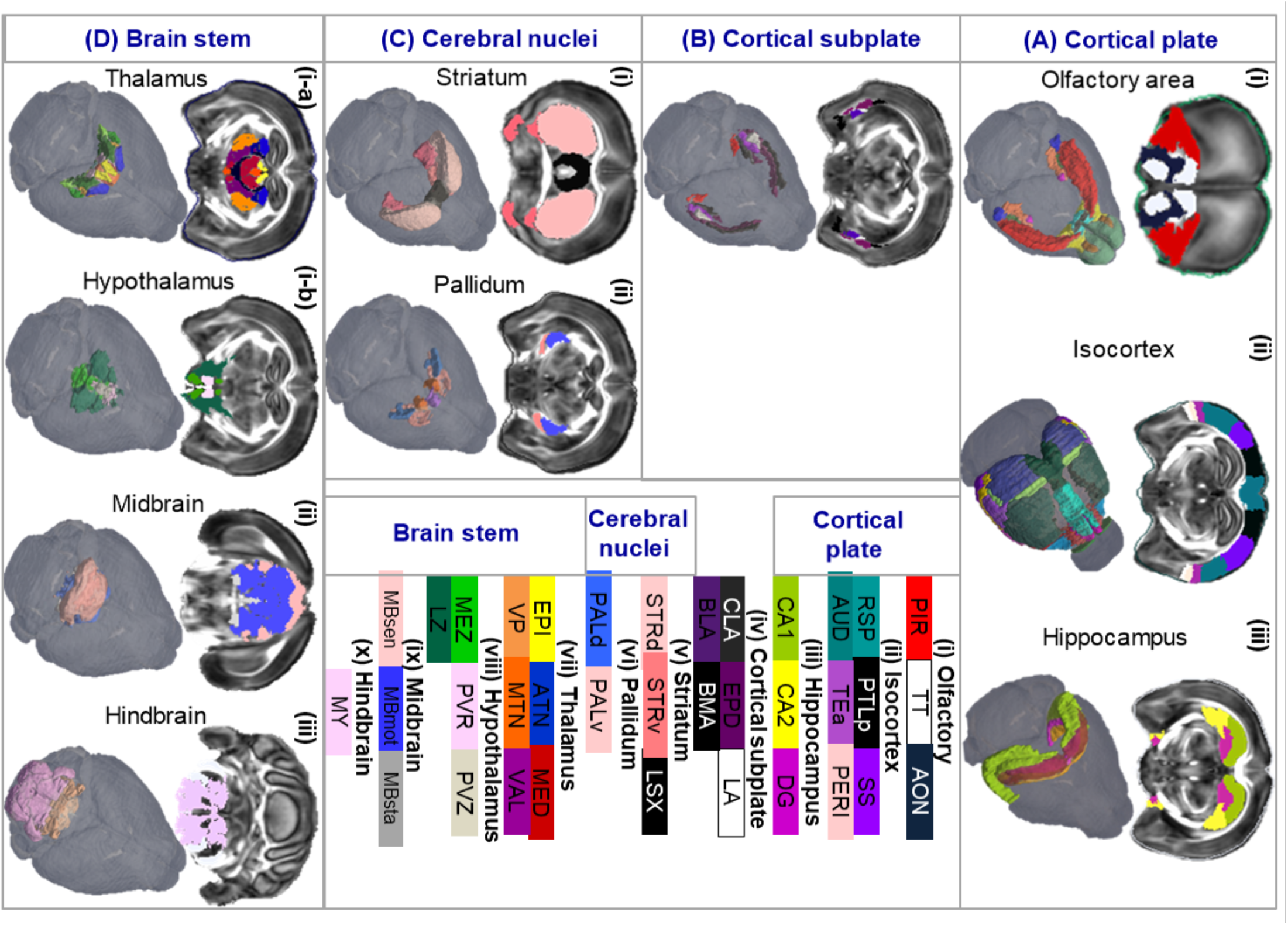
Segmentation tree illustrating the hierarchy of the atlas: A) Cortical plate is partitioned into olfactory, isocortex, and hippocampus regions. B) Cortical subplate/amygdala) and its sub-regions. C) Cerebral nuclei atlas containing the striatum and pallidum. D) Brain stem including the thalamus, hypothalamus, midbrain and hindbrain. Abbreviations of the anatomical structures are same as defined in the AMBA (https://mouse.brain-map.org/static/atlas).

Major cortical regions described in the isocortex atlas (Fig. 4A-ii) were further partitioned into 40 sub-regions in the left and right hemisphere respectively as shown in supplementary Fig. S2 A. For example, somatosensory area (SS), including both primary and secondary SS regions in Fig. 4A-ii, was separated into 7 parts in the left and right hemisphere (Supplementary fig. S2 A: primary somatosensory area – nose (SSp-n), mouth (SSp-m), upper limb (SSp-ul), lower limb (SSp-ll), barrel field (SSp-bfd), trunk (SSp-tr), and secondary somatosensory area (SSs)). Similarly, the hippocampal region was separated into 9 subregions: CA1, CA2, CA3, DG, entorhinal area (ENT), parasubiculum (PAR), postsubiculum (POST), presubiculum (PRE), and subiculum (SUB) (Supplementary fig. S2 B). Additionally, we imported 219 cortical layer labels from AMBA into our dMRI-based atlas and formed a comprehensive bi-lateral atlas of the mouse cortical layers (Supplementary fig. S2 C-D).

### 3.2 Comparison with MEMRI data

As the structural labels were imported into the template space (DWI and FA), based on the tissue contrasts in the AMBA reference images (autofluorescence) and dMRI (FA and DWI, mostly linked to white matter), it is not clear whether boundaries of GM structural labels were mapped consistently as gray matter structures (e.g., cortical area and thalamic nuclei) often lack definitive contrasts in mouse MRI at 7T. Since *in vivo* MEMRI images show T1-enhancement in the hippocampal subregions and in many thalamic nuclei (Szulc et al., 2015), acquired datasets were mapped to the atlas following acquisition from a separate cohort to qualitatively examine the spatial agreement of several gray matter structural labels. Fig. 5A shows three axial slices (top to bottom) of an average of MEMRI images (n = 6) overlaid with structural labels mapped using affine transformations. Regions of several thalamic nuclei from the atlas (e.g., epithalamus (EPI), reticular nucleus (RT), ventral posterior complex of the thalamus (VP), dorsal part of the lateral geniculate complex (LGd), geniculate group, ventral thalamus (GENv), lateral group of the dorsal thalamus (LAT) can be distinguished in the MEMRI data based on their intensity levels (Fig. 5B).

**Fig. 5:**
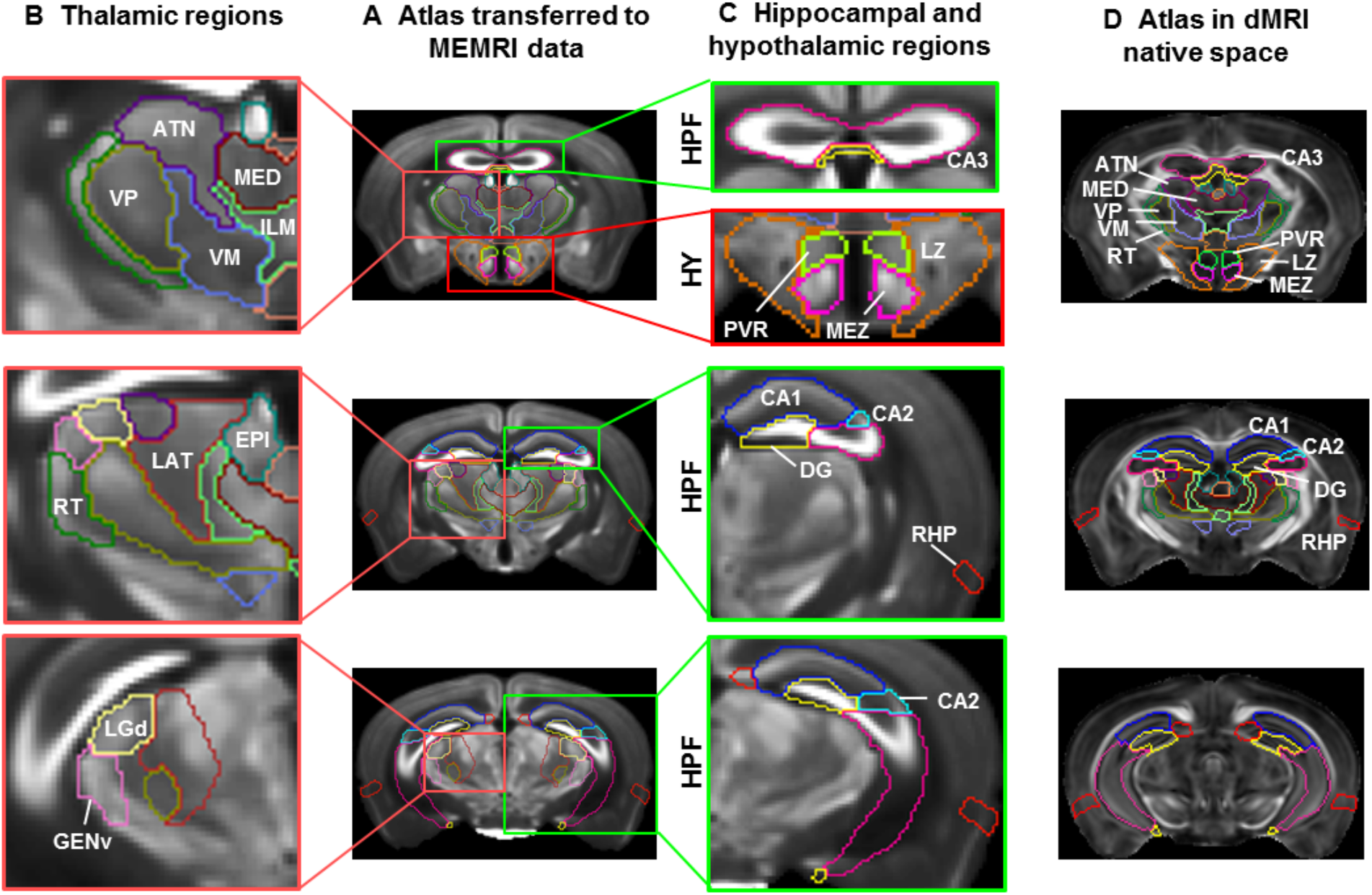
Comparisons of the dMRI-based atlas with MEMRI data: A) Representative axial slices of an in vivo MEMRI data overlaid with affine-aligned structural labels in the hippocampus (HPF, green rectangle), thalamus (pink rectangle) and hypothalamus (HY, red rectangle). B) Magnified view of the thalamic region showing several nuclei of the thalamus–reticular nucleus (RT), Ventral posterior complex of the thalamus (VP), dorsal part of the lateral geniculate complex (LGd), geniculate group, ventral thalamus (GENv), and lateral group of the dorsal thalamus (LAT). C) Magnified view of the hippocampal and hypothalamic regions, showing highlighted neuronal layers in the CA1, CA2, CA3, and dentate gyrus (DG) of the hippocampus and hypothalamic region highlighting paraventricular, lateral, and medial zone (PVR, LZ, and MEZ respectively). D) The structural labels used in B and C overlaid on FA images in the dMRI space for comparison.

Similarly, regions of several hypothalamic nuclei from the atlas (e.g., periventricular region (PVR), medial zone (MEZ), and lateral zone (LZ)) also matched the MEMRI results (Fig. 5C – upper panel). Magnified view of the hippocampal regions (Fig. 5C) showed enhanced neuronal layers that fitted the defined boundaries of hippocampal regions (e.g., CA1, CA2, and DG in Fig. 5C). The ventral part of the hippocampus region in the atlas did not align well with the in vivo MEMRI data (Fig. 5C – lower panel), due to changes in brain morphology after death or fixation and may be corrected by nonlinear brain mappings.

### 3.3 Mapping mouse brain thalamocortical connectivity

Ipsi-lateral thalamocortical connections between cortical seed regions and target regions in the thalamus were reconstructed using dMRI (Fig. 6A and C) and compared with corresponding tracer maps (Fig. 6B, injections sites identical to the seed regions) for their spatial overlaps (Fig. 6D-E).

**Fig. 6:**
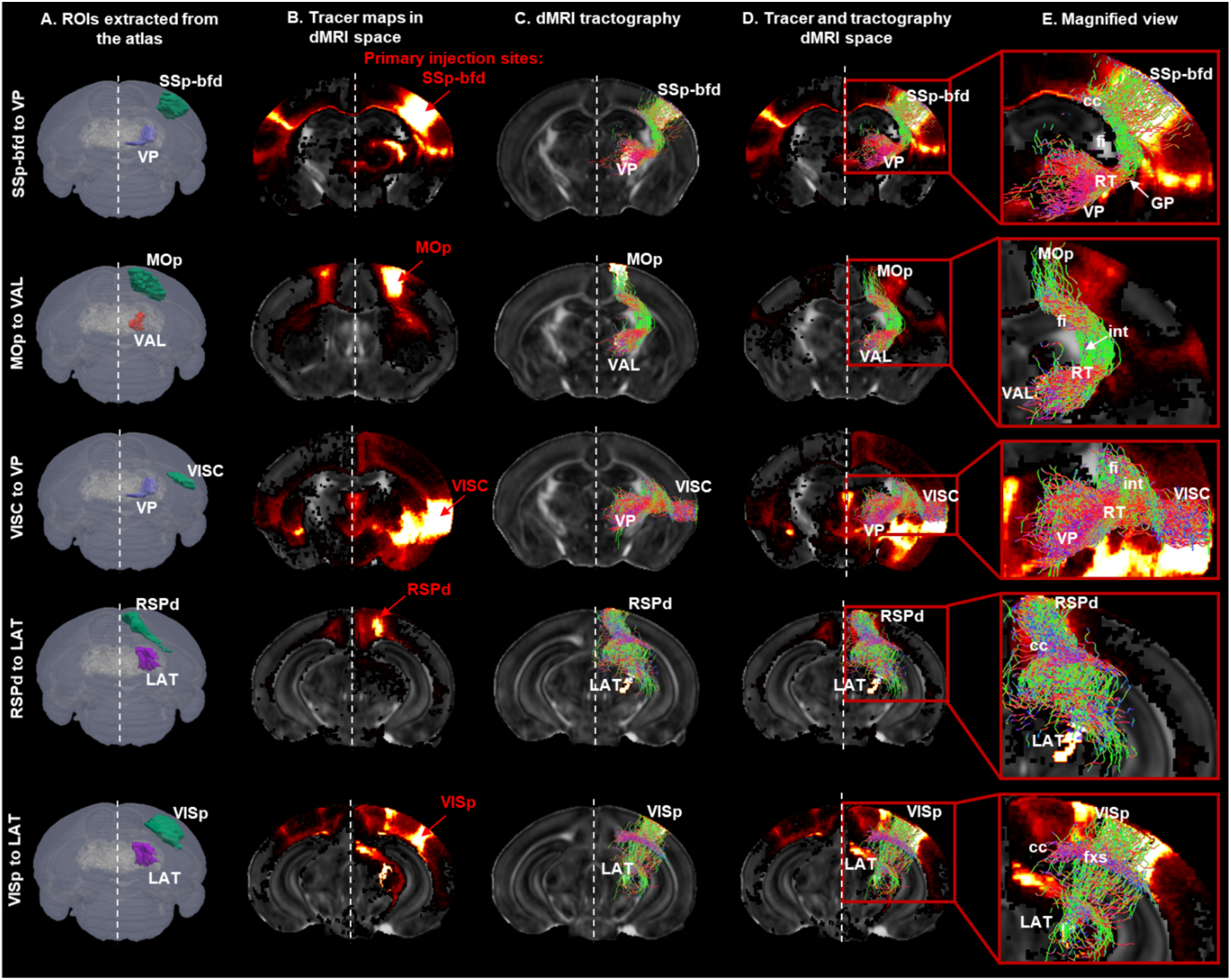
Validating dMRI tractography using tracer maps from AMBCA: A) Ipsi-lateral cortical and thalamic ROIs extracted from the dMRI-based atlas used for fiber tractography: (from top to bottom) somatosensory barrel-field (SSp-bfd), ventral posterior complex of the thalamus (VP), primary motor area (MOp), ventral anterior-lateral complex of the thalamus (VAL), primary visual area (VISp), lateral group of the dorsal thalamus (LAT), retrosplenial area – dorsal part, and visceral area (VISC). B) Representative tracer maps for the selected injection sites (from top to bottom: SSp-bfd, MOp, VISp, RSPd, and VISC) from the AMBCA transferred to the dMRI space overlaid on FA image. C) Ipsilateral fiber projections from the cortical seed regions to the thalamic target regions derived via dMRI tractography: (from top to bottom) SSp-bfd to VP, MOp to VAL, VISp to LAT, RSPd to LAT, and VISC to VP. D) Tracer maps and fiber tractography overlaid onto FA image in the dMRI space. E) Magnified view showing identical patterns of axonal projections as tracer maps.

The tractography results produced a binary 14×12 thalamocortical connectivity matrix. Fig. 7 shows the 14×12 thalamocortical connectivity matrices derived from the tracer data in AMBCA – ground truth (Fig. 7A) and from dMRI tractography under six conditions listed in table 2 (Fig. 7B-C). A detailed description of the connectome construction is provided in the supplementary material. The matrix generated under condition 1 (b = 2,000 s/mm^2^ and 30 diffusion directions using NCT) served as the baseline. Increasing the diffusion weighting to b=5,000 s/mm^2^ (condition 2) recovered several FN connections (blue dots in Fig. 7B) but also introduced several FP connections (black dots in Fig. 7B). Increasing the number of diffusion direction to 60 (condition 3) improved the recovery of FNs without inducing any further FPs. Based on all 10 subjects, condition 3 improved the TP rate from 67.79% to 92.37% (p < 0.0001) and reduced the FN rate from 32.2% to 7.6% (p < 0.0001) over condition 1 but at the cost of an increased FP rate (31.57% to 42.1%, p = 0.0169) (Fig. 7D top panel).

**Fig. 7:**
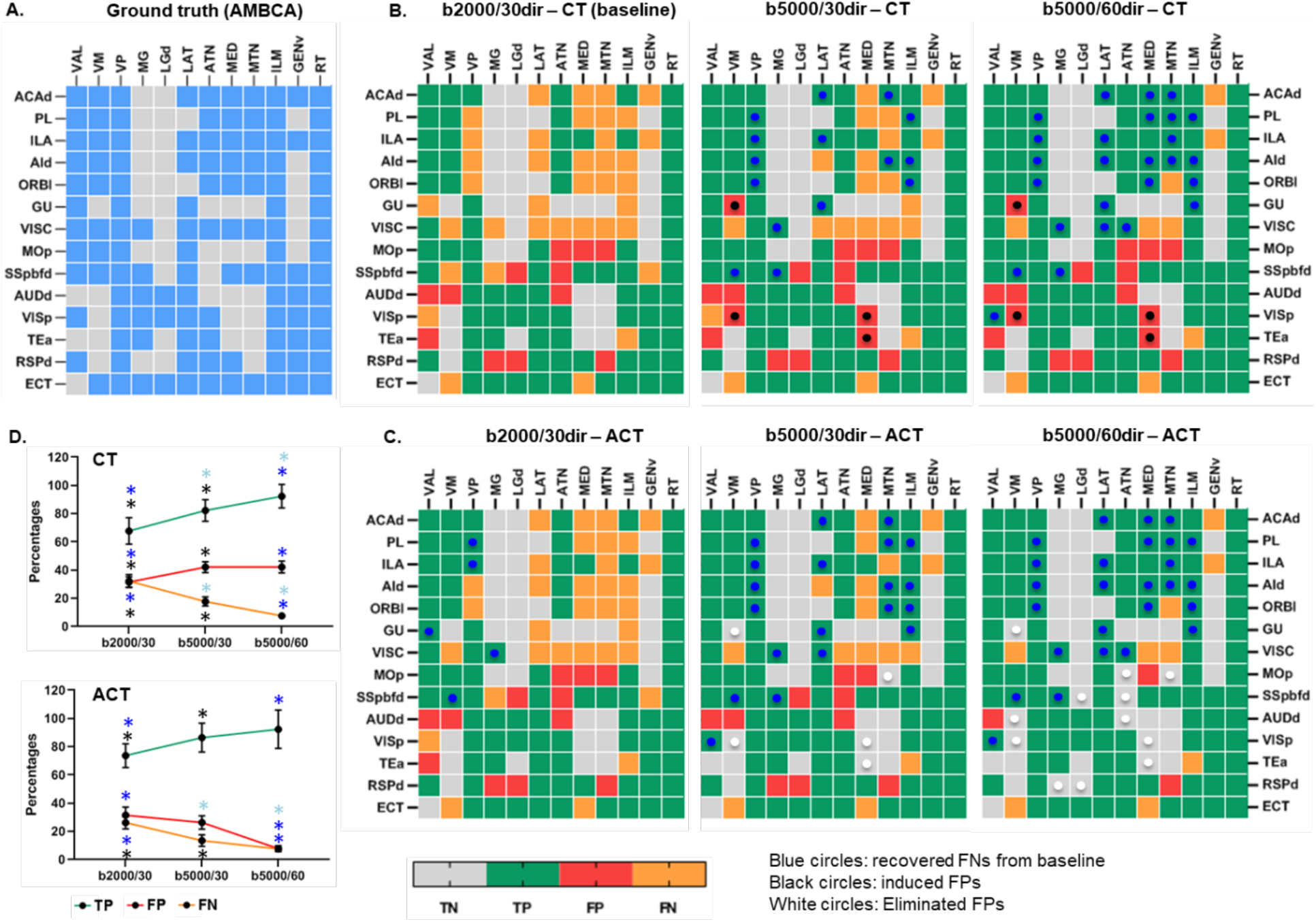
Thalamocortical connectome matrix: A) Generated from Allen Mouse Brain Connectivity Atlas (AMBCA) – ground truth. B) Using conventional tractography (NCT) with different b-values and diffusion directions. C) Using anatomically constrained tractography (ACT).

Compared to the baseline, ACT under condition 4 reduced FN detections by 6% (blue dots) but did not eliminate FPs. Condition 5 on the other hand, increased the detection of TPs (by 18.6%) as compared with the baseline and further removed the FPs by 15.8% that were induced with higher b-values or diffusion directions using NCT (conditions 3 and 4). Notably, condition 6, i.e., b=5,000 s/mm^2^ and 60 diffusion directions using ACT, produced the lowest FP rates (7.89%), with a 92.37% TP rate and thus outperforms all other conditions (Two-way ANOVA with Tukey’s multiple comparisons test, α = 0.05). The summary of the findings from all six condition is summarized in Table 4.

**Table 4:** TP, FP, and FN detection rates for each condition in Table 2.

| Conditions | 1: b=2000,<br>30 dirs, NCT | 2: b=5000,<br>30 dirs, NCT | 3: b=5000,<br>60 dirs, NCT | 4: b=2000,<br>30 dirs, ACT | 5: b=5000,<br>30 dirs, ACT | 6: b=5000,<br>60 dirs, ACT |
| --- | --- | --- | --- | --- | --- | --- |
| TP rate<br>(%) | 67.8 | 82.2 | 92.4 | 73.7 | 86.4 | 92.4 |
| FN rate<br>(%) | 32.2 | 17.8 | 7.6 | 26.3 | 13.6 | 7.6 |
| FP rate<br>(%) | 31.6 | 42.1 | 42.1 | 31.6 | 26.3 | 7.9 |

Visual comparisons of the NCT and ACT results showed ACT streamlines were more likely to stay in WM than NCT streamlines (Supplementary Fig. S3A). In addition, compared to NCT, the ACT results showed marked reductions in ill-defined fiber termination points, such as, fibers that ended in the WM or CSF regions (Supplementary Fig. S3B). This difference was expected as ACT checked termination conditions, which may help ACT to identify more TP connections while minimizing the FP detections for the same imaging setups.

### 3.4 Spatial overlap between tractography and tracer

While dMRI tractography can find streamlines passing through a pair of cortical and thalamic regions, the trajectories of tractography streamlines, in some cases, differed from those of the tracer results (Fig. 8). Fig. 8 illustrates some examples of the open-end fiber projection maps derived from the six conditions and quantification of the spatial overlaps with corresponding tracer data. We observed that both imaging parameters and tractography methods had effects on the level of spatial overlap between the tracer and tractography results, and the effects differed among cortico-thalamic connections. For example, both imaging parameters and tractography methods had significant effects (Two-way ANOVA with Tukey’s multiple comparisons test, α = 0.05) on the SSp-bfd projection map (Fig. 8A). However, only diffusion imaging parameters had a significant effect on the DICE score for dorsal auditory (AUDd) projection map (Fig. 8B), whereas, for the primary visual area tractography method ACT performed better than the NCT method (Fig. 8C).

**Fig. 8:**
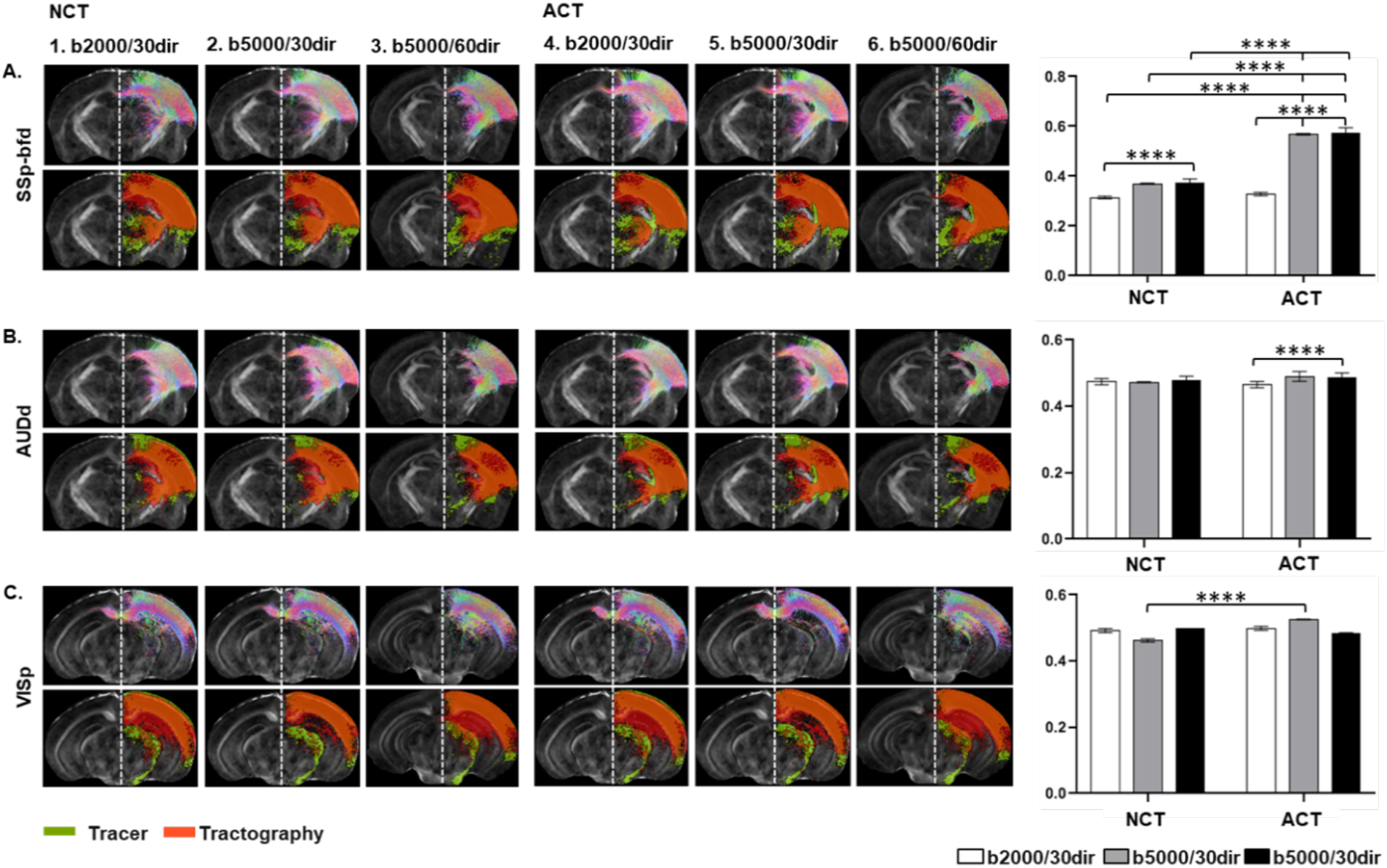
Axial slices showing open-end fiber tractography (upper panel) and corresponding tracer maps and tract density images using six conditions (table 2) overlaid on subject’s FA image: A) Somatosensory barrel field (SSp-bfd, B) Auditory cortex - dorsal area (AUDd) and C) Visual cortex – primary area (VISp).

As each tracer data may contain multiple connections within the thalamocortical network, it is difficult to segregate the tracts and form ground truth for node-to-node tractography comparisons. Instead, we compared the portion of each thalamic nuclei occupied by tracer (valid termination sites) and tractography streamlines from comparable cortical regions. Fig. 9 shows the DICE scores in detected TP connections in the thalamocortical circuitry under different conditions (1, 2, 3, 5, and 6). Green, yellow, and blue cells represent good, moderate, and poor spatial agreements in termination between the tractography and tracer data between the corresponding cortical and thalamic nodes. From cortical node-to-thalamus quantification, we identified three groups of connections (SSp-bfd to TH, MOp to TH, and VISp to TH) that showed 80% or more similarities with the corresponding tracer maps. Another three groups of connections (AUDd to TH, RSPd to TH, and TEa to TH) showed 60% to 80% similarities, and the remaining 8 group of connections showed poor agreement (below 60%) with the tracer maps. Three representative images showing quantification of the DICE scores for different tracts from three different levels have been shown in the supplementary Fig. S4.

**Fig. 9.**
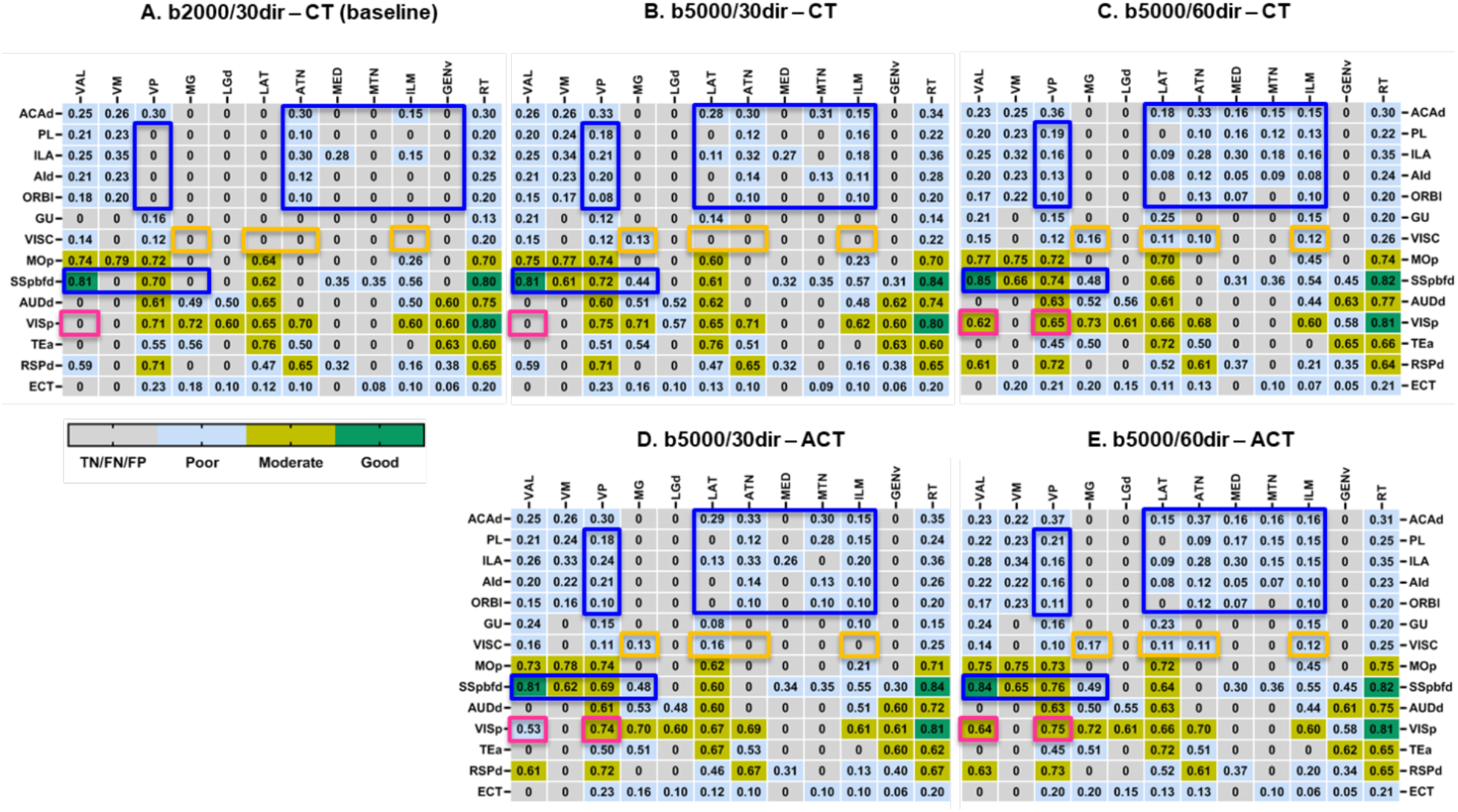
Tract specific DICE scores of thalamic termination points in the thalamocortical network computed from six conditions in Table 2. For each condition, DICE scores measuring the spatial agreement in thalamic termination points for each TP connections are displayed and classified. Dice scores of connections within the blue rectangles improved with both higher b-values and more diffusion directions, and dice scores of connections within the orange rectangles improved with either higher b-values or more diffusion directions. Dice scores of connections within the magenta rectangles improved with switching from NCT to ACT.

While the spatial agreements for thalamic termination point remained poor for most connections, we observed that the imaging parameters and tractography methods have tracts specific effects within the thalamocortical network. For example, in addition to the tractography techniques, higher b-values and diffusion directions can significantly improve the detection of TP connections along the prefrontal-thalamic and the sensory-thalamic pathways (Fig. 9 A-E, blue boxes). Changing only the b-values and diffusion directions on the other hand significantly impact the interactions between visceral to the polymodal association cortex related thalamic nuclei (LAT, ATN, ILM) and medial geniculate complex (MG) of the dorsal thalamus (MG/LGd) (Fig. 9 A-E, indicated by orange boxes), whereas tractography method shows significant effect on the visual to the ventral group of dorsal thalamic pathway (Fig. 9 A-E, magenta boxes).

## Discussion

Here, we reported an improved dMRI-based atlas of the adult C57BL/6J mouse forebrain integrated with AMBA structural labels. We demonstrated utility in reconstructing the mouse thalamocortical connectivity from dMRI data. The long-term goal of this atlas is accurate assessment of structural connectome of the mouse brain that would be useful to extend in various mouse models.

In accordance with the structural hierarchy defined in the AMBA, our dMRI-based atlas includes structural labels of the major GM regions at different cortical and subcortical levels. However, the boundaries of GM subregions can be ambiguous due to limited MRI resolution and nonspecific contrast patterns. For example, our isocortex atlas includes 17 cortical regions such as FRP, MO, SS, etc., which was additionally segmented into dorsal, ventral, anterior, and posterior regions following the AMBA, resulting a detailed atlas of the mouse cortex containing 80 structures segregated equally in both hemispheres. These regions, originally defined in the AMBA, were imported using a mapping that was computed based on segmentation of major brain compartments, not based on specific contrasts in dMRI signals. For the same reason, our atlas lacks in-depth classification of the subcortical layers and nuclei as labeled in the AMBA. Even though the comparison using the MEMRI data demonstrated good agreement in several hippocampal and thalamic regions in normal mouse brains, the accuracy of other regions has not been evaluated due to lack of good tissue contrasts that match histological methods. Applying the atlas to mutant mouse brains with altered cortical regions and cytoarchitecture or severe deformation, may result in inaccurate delineation of cortical and other GM regions.

Although major WM pathways can be reliably reconstructed using dMRI tractography, taking the advantage of the atlas, we demonstrated the possibility of seeking neural connections within largely GM structures. We focused on the thalamocortical network because it is a major pathways conserved across multiple species (Garel and Rubenstein, 2004; Leyva-Díaz and López-Bendito, 2013; Rakic, 1975), and contribute in fundamental brain functions (Bagshaw et al., 2014; Sarnthein et al., 2005; Seeley et al., 2007). Understanding the extent to which the reconstructed pathways reflect realistic trajectories is important for future use of dMRI tractography to study the thalamocortical pathways or similar pathways in mouse models. Although we used post mortem mouse brains in this study as it avoids most of the sources known to degrade *in vivo* DW-MRI, such as low SNR, and motion artefacts (Le Bihan et al., 2006; Lori et al., 2002), we have previously demonstrated the feasibility to acquire high-resolution dMRI data from live mouse brains (Wu et al., 2014).

With dMRI datasets and AMBCA tract tracing data coregistered into the template space, we were able to perform a thorough investigation of individual node-to-node connections in the thalamocortical network. Looking at the connectivity matrix, results obtained from the baseline showed good consistency with the ground truth (Fig. 7A-B), albeit FPs and FNs were observed in several connections. FP connections (red cells in Fig. 7B) were detected in five out of fourteen cortical nodes (MOP, SSp-bfd, AUDd, Tea, and RSPd) to the thalamic network, and FN connections (orange cells in Fig. 7B) were predominantly associated with several thalamic regions (e.g., MED, MTN, ILM, LAT, and VP). Identification of FPs and FNs indicates the gap between tractography results and tracer ground truth for connections passing through gray matter regions. These findings not only suggest the importance of the validation of tractography maps by AMBCA, but also highlight the necessity of optimizing the tractography method to control the false positives or negatives.

We addressed this issue by applying additional constraints based on the anatomical knowledge of the brain tissues while generating the streamlines that infer more realistic microstructural details in relation to the inherent biology. Despite the NCT-based thalamocortical tracts seeming reasonable, we observed significant improvement in the overall connectome using ACT. In addition, varying the b-values and diffusion directions, we identified suitable imaging parameters (b-value/ number of diffusion directions) that can be used for more reliable estimation of the thalamocortical tracts. For example, increasing the imaging parameters from baseline conditions improved the rate of TP detections with trivial improvement in FP detections. However, altering both the imaging and tractography methods yielded significant improvement in TP detections with minimal FP and FN connections. These findings suggest that without anatomical priors, the NCT approach accepts biologically implausible fiber trajectories as valid termination points. In contrast, the ACT approach allows tracing the continuation of streamlines through GM regions using the acceptance/rejection criteria. Nevertheless, we could not detect all TP connections within the thalamocortical network, which still suffer from FP biases and may be likely due to the differences in FOD profiles between tractography and tracer maps that warrant further investigations.

When we further examined the spatial agreements between tractography and tracer results for each node-to-node connection, only a small number of connections showed good to moderate agreements between the two (Fig. 9). For example, a connection was found between VISC and VAL, but the trajectories of tractography and tracer results are quite different (Fig. 8), resulting in low DICE score. The evidence demonstrates the limitation of dMRI tractography in reconstructing structural connections through mouse brain GM, which is not surprising given the complex microstructural organization often found in GM regions while it is difficult to separate axons from cell bodies and dendrites solely based on dMRI signals. With higher GM to WM compartment ratio in the mouse brain than in the human brain, this limitation may pose a big challenge for studies that use dMRI tractography to examine structural connectivity in the mouse brain.

Acquiring high resolution MR data might be one of the solutions as it will reduce partial volume effect, so that smaller axon bundles can be more easily separated from surrounding tissues, but at an expense of longer data acquisition time (Anderson et al., 2020; Crater et al., 2022). Over the past decade, the compressed sensing (CS) technique has been widely used to speed up the MR data acquisition by under-sampling the *q*-space or the *k*-*space* (Aranda et al., 2015; Daducci et al., 2015; Menzel et al., 2011; Tobisch et al., 2018). Recently, Wang et al. demonstrated the acquisition of 50 μm dMRI datasets of the mouse brain using CS in the k-space (Wang et al., 2020b), and we have reported a joint q-space and k-space CS method that can reconstruct *ex vivo* mouse brain dMRI data with acceleration factors up to 8 (Zhang et al., 2020). Another potential solution is the development of more advanced dMRI acquisition and tractography techniques. For example, the recent reported b-tensor technique (Cottaar et al., 2020) can potentially separate axons from cell bodies by introducing spherical diffusion encoding.

## Conclusion

In conclusion, we have developed an atlas based on dMRI with structural labels imported from the AMBA that can serve as a requisite template for cross-examination of potential disrupted connections/volumetric changes in the mouse brain. Furthermore, comparing dMRI tractography results with AMBCA tracer results revealed significant differences in the trajectories of mouse thalamocortical tracts, indicating the need for further development of dMRI tractography for accurate characterization of mouse brain structural connectivity.

## Acknowledgment

We would like to thank the members of the labs of Drs. Jiangyang Zhang, Daniel Turnbull, Youssef Z. Wadghiri and the team of the preclinical imaging core at the NYU Grossman School of Medicine, for their support and feedback during the evolution of our work.

