## Supplementary Materials for "Towards reliable reconstruction of the mouse brain thalamocortical connectivity using diffusion MRI"

**Description of the mouse brain structures:**

**Whole brain atlas:** Mouse whole brain (Cereberum) atlas (fig. 4a) includes 7 major GM structures such as olfactory area, isocortex, hippocampus, striatum, interbrain, midbrain, and cerebellum. Whole brain atlas was further segmented according to the Allen Mouse Brain Atlas (AMBA) hierarchy.

a) **Cortical plate:** We created three distinct atlases from the mouse brain cortical plate region (fig. 5A):
   (i) **Olfactory area (OLF) atlas:** OLF atlas (fig. 5A - i) is composed of 10 regions, such as, main olfactory bulb (MOB), accessory olfactory bulb (AOB), anterior olfactory nucleus (AON), taenia tecta (TT), dorsal peduncular area (DP), piriform area (PIR), ucleus of the lateral olfactory tract (NLOT), cortical amygdala area (COA), piriform amygdala area (PAA), and post-piriform transition area (TR).
   (ii) **Isocortex (CTX) atlas:** Isocortex atlas (fig. 5A - ii) contains 17 cortical regions: frontal pole (FRP), somatomotor area (MO), somatosensory area (SS), gustatory area (GU), visceral area (VISC), auditory area (AUD), visual area (VIS), anterior cingulate area (ACA), prelimbic area (PIR), infralimbic area (ILA), orbital area (ORB), agranular insular area (AI), retrosplenial area (RSP), posterior parietal association area (PTLp), temporal association area (TEa), perirhinal area (PERI), ectorhinal area (ECT).
   (iii) **Hippocampus (HPF) atlas:** HPF atlas (fig. 5A - iii) is made of 5 major hippocampal regions: ammon's horn - field CA1, field CA2, field CA3, dentate gyrus (DG) and retrohippocampal region (RHP). RHP is a collective region combining entorhinal area (ENT), parasubiculum (PAR), postsubiculum (POST), presubiculum (PRE), and subiculum (SUB).

b) **Cortical subplate (CTXsp) atlas:** Atlas of the cortical subplate (fig. 5B), also known as amygdala, embraces major amygdalar nucleuses, such as, endopirirform nucleus (EP), lateral amygdalar nucleus (LA), basolateral amygdalar nucleus (BLA), basomedial amygdalar nucleus (BMA), posterior amygdalar nucleus (PA), as well as claustrum (CLA).

**c)** Mouse brain cerebral nuclei (fig. 5C) is segmented into two structures:

(i) **Striatum (STR):** Striatal atlas (fig. 5C - i) has 4 discrete structures: dorsal (caudate putamen) and ventral (nucleus accumbens) regions of the STR, plus the lateral septal complex (LSX) and striatum like amygdalar nuclei (sAMY).

(ii) **Pallidum (PAL):** PAL atlas (fig. 5C - ii) contains dorsal (PALd), ventral (PALv), medial (PALm), and caudal (PALc) regions of the pallidum.

**d)** Brain stem area (fig. 5D) is partitioned into 3 structures:

(i) **Interbrain (IB):** Mouse interbrain is divided into thalamus and hypothalamus.

(i-a) **Thalamus (TH) atlas:** Thalamus atlas embraces 14 thalamic nucleis: ventral anterior-lateral complex of the thalamus (VAL), ventral medial nucleus of the thalamus (VM), ventral posterior complex of the thalamus (VP), subparafascicular nucleus (SPF), subparafascicular area (SPA), peripeduncular nucleus (PP), medial geniculate complex (MG), dorsal part of the lateral geniculate complex (LGd), lateral group of the dorsal thalamus (LAT), anterior group of the dorsal thalamus (ATN), medial group of the dorsal thalamus (MED), midline group of the dorsal thalamus (MTN), intralaminar nuclei of the thalamus (ILM), reticular nucleus of the thalamus (RT), geniculate group, ventral thalamus (GENv), epithalamus (EP).

(i-b) **Hypothalamus (HY) atlas:** HY atlas includes 4 hypothalamic zones: periventricular zone (PVZ), PV region (PVR), hypothalamic medial and lateral zone (MEZ and LZ).

(ii) **Midbrain (MB):** MB atlas is comprised of 3 regions related to sensory, motor, and behavioral state named as MBsen, MBmot, and MBsta.

(iii) **Hindbrain (HB):** HB atlas has 3 structures from pons: sensory related (P-sen), motor related (P-mot), behavioral state related (P-sat) and similarly 3 more structures from medulla (MY): MY-sen, MY-mot, and MY-sat.

Generation of the mouse brain thalamocortical structural connectome: Using the Allen Mouse Brain Connectivity Atlas (AMBCA), firstly, we constructed the tracer-based thalamocortical connectivity matrix. Instead of considering the projection densities from a cortical injection site to a voxel in the thalamus, we rather considered whether a projection is existed in that particular voxel or not and marked as 1 (connected) or 0 (not connected) respectively. Subsequent binary matrix served as the ground truth (fig. 9A) for further quantitative analysis.

Next, we constructed dMRI tractography based thalamocortical connectome under different conditions as described in table 2. Two nodes were considered to be connected if one or more streamlines had their endpoints in both nodes. All detected node-to-node connections were marked as ‘1’, otherwise 0 (not connected). This resulted in six 14×12 binarized structural connectivity matrix (fig. 9B-C). For both matrices, cortical nodes were assigned in rows and thalamic nodes in columns. Each cell of the matrix denotes whether there is a connection present in between the thalamocortical nodes or not. Each row represents the connectivity status between the corresponding cortical and thalamic nodes. Gray, green, red, and orange colors represent true negative (TN), true positive (TP), false positive (FP), and false negative (FN) connections.

| Author (s) | Mouse strain(s) | Imaging modalities | Brain regions covered | Number of structures | Resolution (µm³) |
|---|---|---|---|---|---|
| Steadman et al., 2014 | NL3 KI | MR | Cerebellum | 39 | 56 |
| Ullmann et al., 2013 | C57BL/6 | MR | Neocortex | 74 | 15 |
| Richards et al., 2011 | C57BL/6 | MR | Hippocampus | 40 | 30 |
| Sharief et al., 2008 | C57BL/6 And BXD | MR | Whole brain | 33 | 43 |
| Dorr et al., 2008 | C57BL/6 | MR | Whole brain | 62 | 32 |
| Dorr et al., 2007 | CBA | MR and CT | Whole brain & vasculature | 26 | 32 |
| Badea et al., 2007 a & b | C57BL/6 | MR | Whole brain | 33 | 43 |
| Chen et al., 2006 | C57BL/6, 129SI/Svlm, CD1 | MR | Whole brain | 42 | 60 |
| Bock et al., 2006 | cdf/cdf, cdf/+ and +/+ | MR and histology | Cerebrum & cerebellum | 5 | 156 |
| Kovacevic et al., 2005 | 129SI/Svlm | MR | Whole brain | 42 | 60 |
| Ma et al., 2005 | C57BL/6 | MR | Whole brain | 20 | 47 |
| Lee et al., 2005 | C57BL/6 | MR | Whole brain | 13 | 40 |
| Ali et al., 2005 | C57BL/6 | MR | Whole brain | 21 | 90 |
| MacKenzie-Graham et al., 2004 | C57BL/6 | MR and histology | Whole brain | 70 | 60 |
| Johnson et al., 2007 | C57BL/6 | MR | Whole brain | 33 | 43 |

| | | | | | |
|---|---|---|---|---|---|
| Chuang et al., 2012 | C57BL/6 | MR | Developing brain (cortex, hippocampus, cerebellum, and white matter tracts) | | 80 to 125 |
| Szulc et al., 2015 | FVB/N | MR | Developing brain (whole brain) | | 100 |

Table S1: Summary of MR-based mouse brain atlases

| | **dMRI atlas** | **MEMRI** | **Tractography** |
|---|---|---|---|
| Imaging | *Ex vivo* | *In vivo* | *Ex vivo* |
| Mouse strain | C57BL/6 | C57BL/6 | C57BL/6 |
| n | 10 | 6 | 10 |
| Sequence | 3D DW multiple spin-echo | T1-weighted 3D spoiled gradient-echo | 3D DW-GRASE |
| Resolution | 125 $\mu m^3$ | 100 $\mu m^3$ | 100 $\mu m^3$ |
| Scanner | Bruker 11.7T | Bruker 7T | Bruker 7T |

Table S2: Summary of the experiments carried out in the study

| Structure names | 3D structures | Structure names | 3D structures |
|---|---|---|---|
| Primary visual area (VISp) | 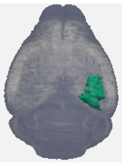 | Retrosplenial area, dorsal part (RSPd) | 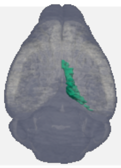 |
| Temporal association area (TEa) | 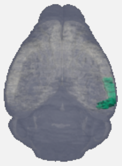 | Anterior cingulate area, dorsal part (ACAd) | 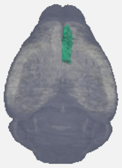 |
| Primary somatosensory area,  barrel-field (SSp-bfd) | 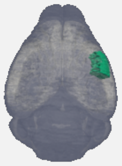 | Prelimbic area (PL) | 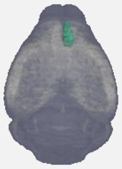 |
| Gustatory area (GU) | 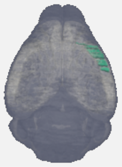 | Infralimbic area (ILA) | 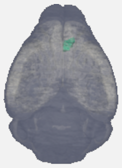 |
| Visceral area (VISC) | 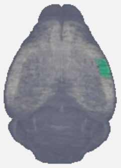 | Orbital area, lateral part (ORBl) | 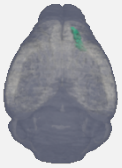 |
| Dorsal auditory area (AUDd) | 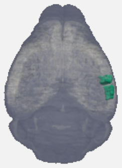 | Agranular insular area, dorsal part (AId) | 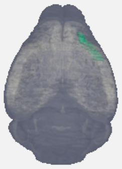 |
| Primary motor area (MOp) | 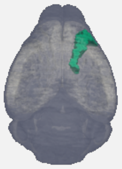 | Ectorhinal area (ECT) | 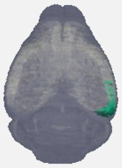 |

Table S3: List of 14 cortical ROIs used for open-end tractography

| Structure names | | 3D structures | Structure names | | 3D structures |
|---|---|---|---|---|---|
| Ventral anterior-lateral complex of the thalamus (VAL) | | 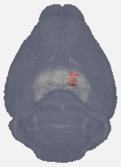 | Anterior group of the dorsal thalamus (ATN) | | 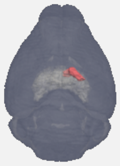 |
| Ventral medial nucleus of the thalamus (VM) | | 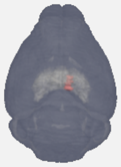 | Medial group of the dorsal thalamus (MED) | | 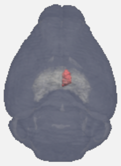 |
| Ventral posterior complex of the thalamus (VP) | | 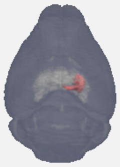 | Midline group of the dorsal thalamus (MTN) | | 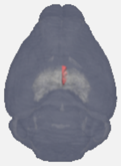 |
| Medial geniculate complex (MG) | | 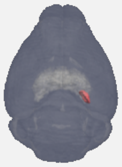 | Intralaminar nuclei of the thalamus (ILM) | | 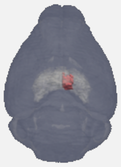 |
| Dorsal part of the lateral geniculate complex (LGd) | | 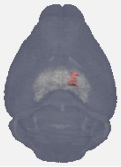 | Geniculate group, ventral thalamus (GENv) | | 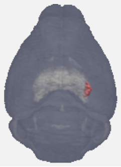 |
| Lateral group of the dorsal thalamus (LAT) | | 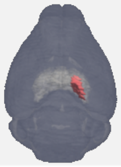 | Reticular nucleus of the thalamus (RT) | | 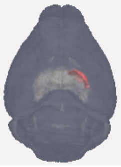 |

Table S4: List of 12 thalamic ROIs used for node-to-node thalamocortical tractography

| Injection sites | Experiment IDs | Reference 3D images from AMBCA |
|---|---|---|
| Primary visual area (VISp) | 309004492 | 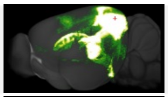 |
| Primary somatosensory area, barrel-field (SSp-bfd) | 112951804 | 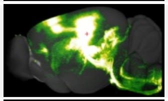 |
| Gustatory area (GU) | 272737914 | 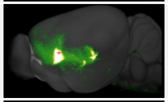 |
| Visceral area (VISC) | 180436360 | 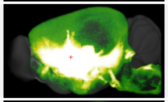 |
| Dorsal auditory area (AUDd) | 158314278 | 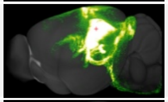 |
| Primary motor area (MOp) | 100141273 | 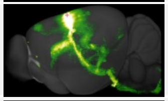 |
| Retrosplenial area, dorsal part | 272916915 | 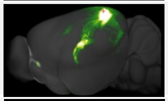 |
| Anterior cingulate area, dorsal part | 159330754 | 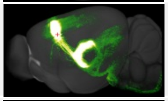 |
| Prelimbic area | 157711748 | 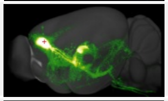 |
| Infralimbic area | 157556400 | 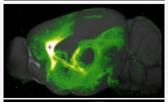 |
| Orbital area, lateral part | 180709230 | 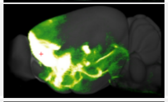 |
| Agranular insular area, dorsal part | 112596790 | 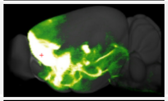 |
| Ectorhinal area | 180435652 | 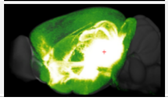 |

Table S5: List of injection sites from the AMBCA (ground truth) used for dMRI tractography validation

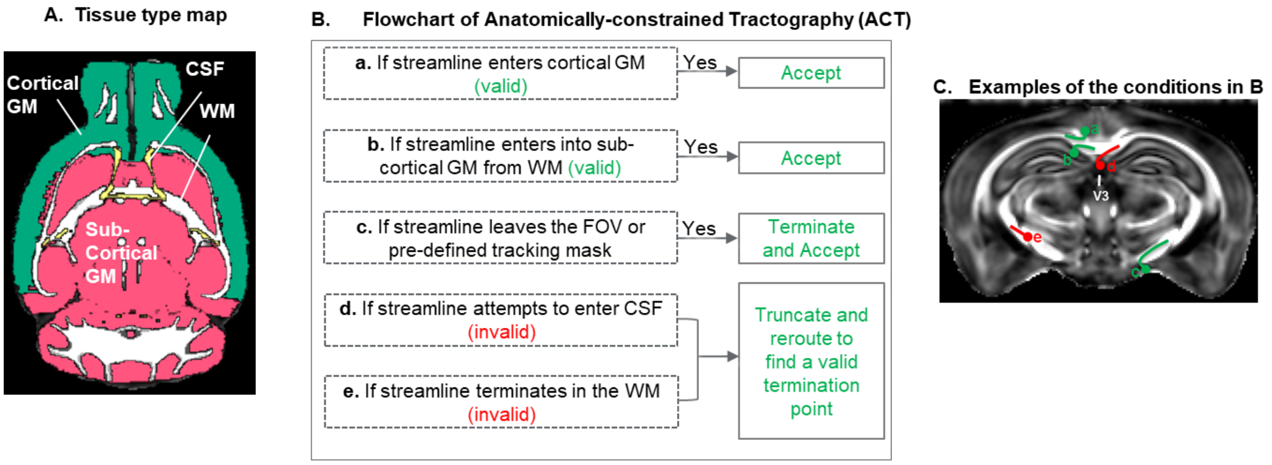


**Supplementary Fig. S1:** Steps in anatomically constrained tractography (ACT). A) Pseudo-colored tissue type map of a mouse brain. B) A flowchart of the streamline termination criteria used by ACT. C) Examples of streamline termination conditions outlined in B).

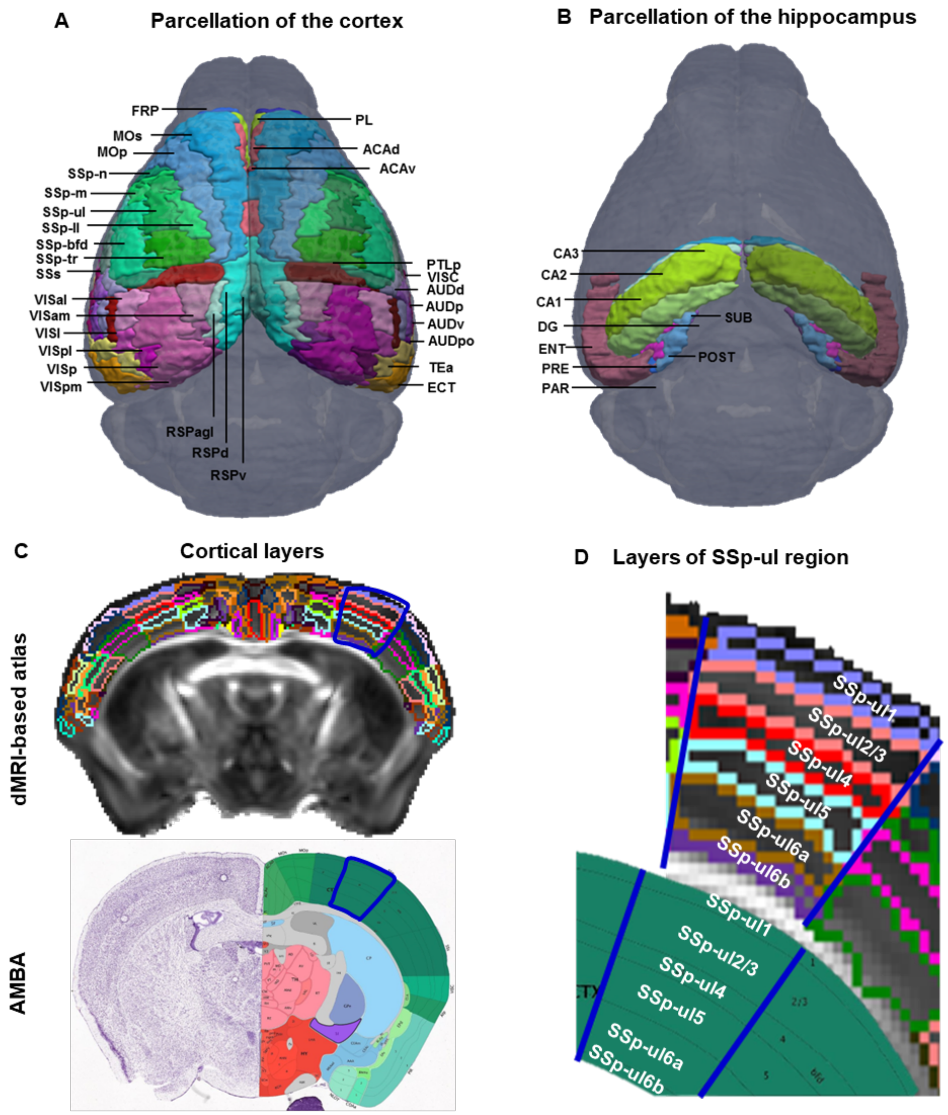


**Supplementary Fig. S2:** Detailed parcellation of the mouse cortex, hippocampus, and cortical layers: Dorsal view of the surface rendering of the parcellations of the cortex (A) and hippocampus (B). C) Mouse brain cortical layers containing 219 structures in AMBA were also imported into the dMRI-based atlas. (D) Magnified view of the six layers of primary somatosensory area, upper limb (SSp-ul) accompanied with the corresponding slice from AMBA (lower panel), showing SSp-ul1 (layer 1), SSp-ul2/3 (layer 2/3), SSp-ul4 (layer 4), SSp-ul5 (layer 5), SSpul-6a (layer 6a) and SSpul-6b (layer 6b). Abbreviations of the anatomical structures are same as defined in the AMBA (https://mouse.brain-map.org/static/atlas).

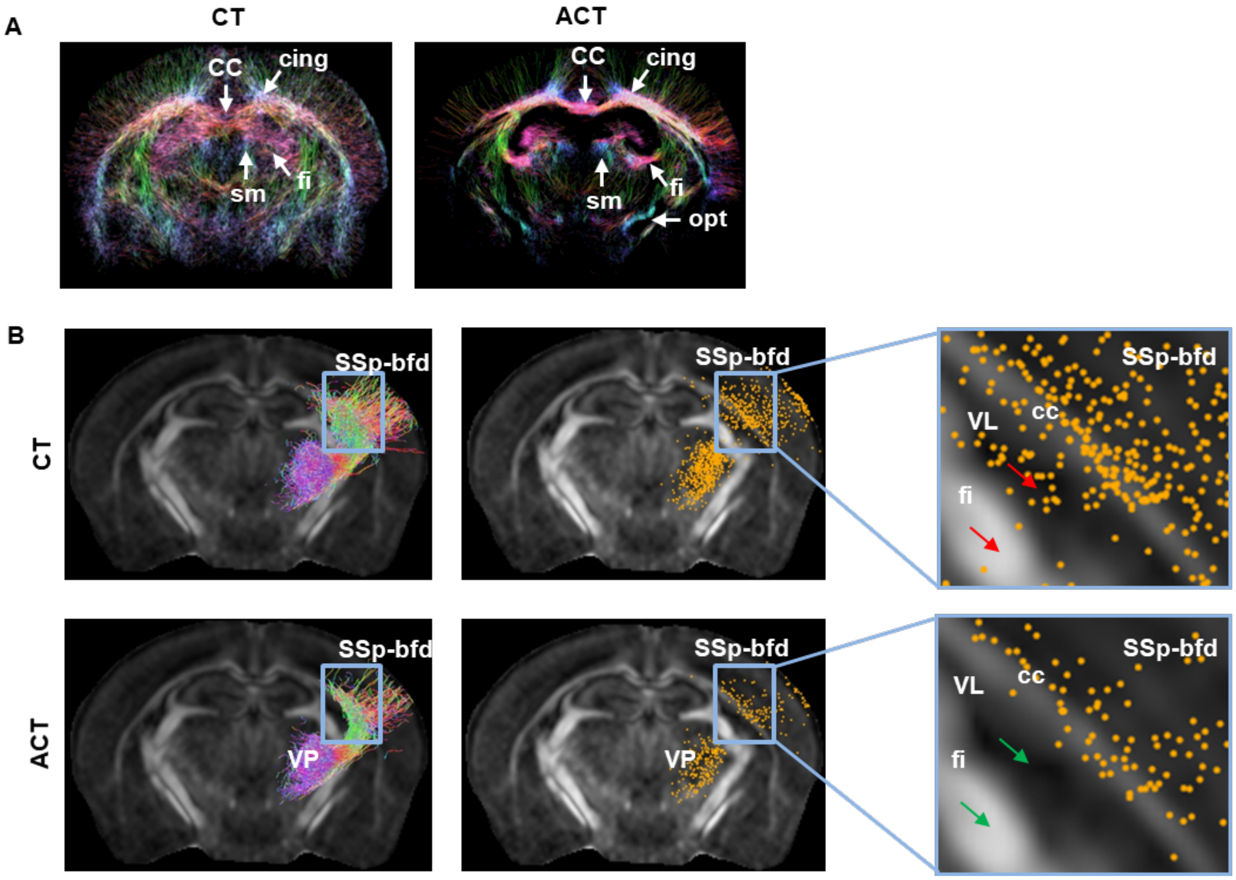


**Supplementary Fig. S3:** Qualitative assessment of the performance of ACT: A) in the reconstruction white matter fiber tracts, B) in reducing the ill-defined termination of the fiber tracts

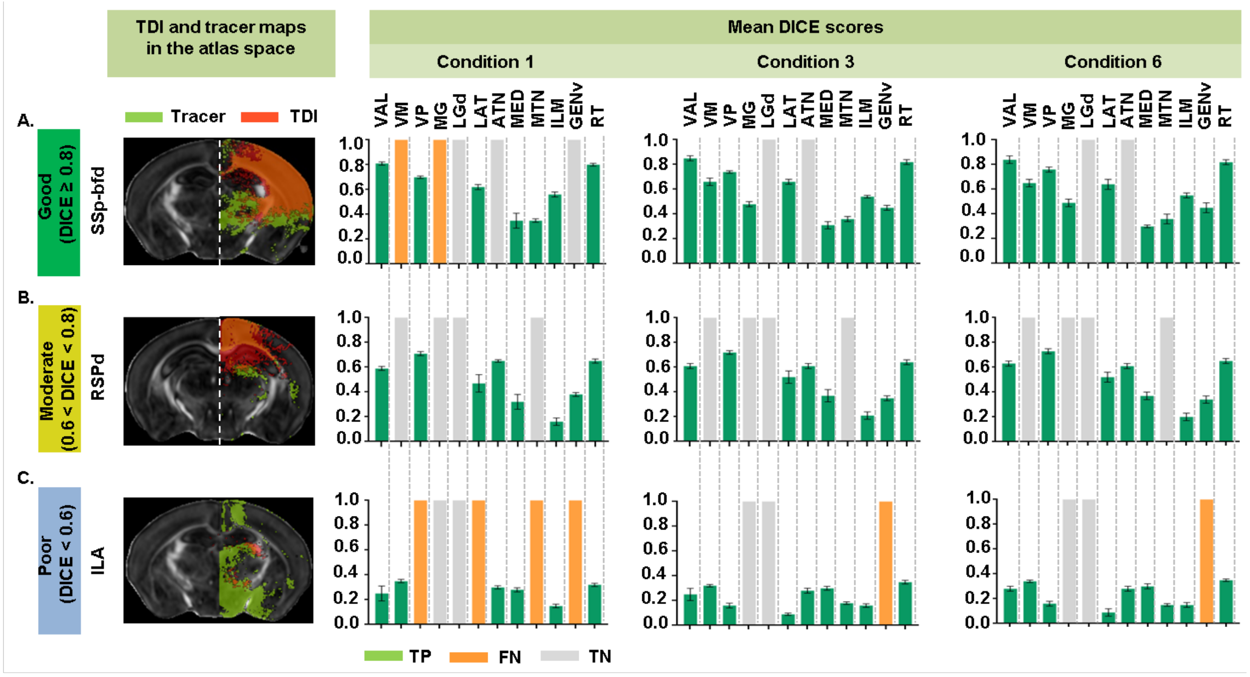


**Supplementary Fig. S4:** Different levels of DICE scores for cortical node-to-thalamus tractography termination points for conditions 1, 3, and 6: Left panel of the figure shows representative brain slices of the TDI and tracer maps of 3 cortical regions, co-registered into the atlas space that show A. good (SSp-bfd), B. moderate (RSPd) or C. poor (ILA) agreement with the ground truth. Right panel of the figure shows quantification of the DICE scores for each tract. Green, orange, and gray bars represent the TP, FN, and TN connections respectively.
